# Healthspan pathway maps in *C. elegans* and humans highlight transcription, proliferation/biosynthesis and lipids

**DOI:** 10.1101/355131

**Authors:** Steffen Möller, Nadine Saul, Alan A. Cohen, Rüdiger Köhling, Sina Sender, Hugo Murua Escobar, Christian Junghanss, Francesca Cirulli, Alessandra Berry, Peter Antal, Priit Adler, Jaak Vilo, Michele Boiani, Ludger Jansen, Dirk Repsilber, Hans Jörgen Grabe, Stephan Struckmann, Israel Barrantes, Mohamed Hamed, Brecht Wouters, Liliane Schoofs, Walter Luyten, Georg Fuellen

**Affiliations:** Rostock University Medical Center, Institute for Biostatistics and Informatics in Medicine and Aging Research (IBIMA), Rostock, Germany; Humboldt-University of Berlin, Institute of Biology, Berlin, Germany; Department of Family Medicine, University of Sherbrooke, Sherbrooke, Canada; Rostock University Medical Center, Institute for Physiology, Rostock, Germany; Rostock University Medical Center, Klinik für Hämatologie, Onkologie und Palliativmedizin, Rostock, Germany; Center for Behavioral Sciences and Mental Health, Istituto Superiore di Sanità, Italy; Budapest University of Technology and Economics, Budapest, Hungary; Abiomics Europe Ltd., Hungary; Institute of Computer Science, BIIT research group, University of Tartu, Tartu, Estonia; Max-Planck Institute for Molecular Biomedicine, Münster, Germany; Ruhr-Universität Bochum, Philosophisch-Theologische Grenzfragen, Bochum, Germany; Universität Rostock, Institut für Philosophie, Rostock, Germany; School of Medical Sciences, University of Örebro, Sweden; University Medicine Greifswald, Klinik und Poliklinik für Psychiatrie und Psychotherapie, Greifswald, Germany; University Medicine Greifswald, Institute for Community Medicine, Greifswald, Germany; KU Leuven, Department of Biology, Leuven, Belgium; KU Leuven, Department of Pharmaceutical & Pharmacological Sciences, Leuven, Belgium

## Abstract

The molecular basis of aging and of aging-associated diseases is being unraveled at an increasing pace. An extended healthspan, and not merely an extension of lifespan, has become the aim of medical practice. However, a precise definition of health and healthspan is not straightforward, and the causal molecular basis of health “per se” is largely unknown. Here, we define health based on the absence of diseases and dysfunctions. Based on an extensive review of the literature, in particular for humans and *C. elegans*, we compile a list of features of health and of the genes associated with them. Clusters of these genes based on molecular interaction data give rise to maps of healthspan pathways for humans, featuring the themes *transcription initiation*, *proliferation* and *cholesterol/lipid processing*, and for *C. elegans*, featuring the themes *immune response*, *mitochondrion* and *biosynthesis* based on genetic and compound intervention data, and *lipids, biosynthesis* and *transcription* based on WormBase compound intervention data. Overlaying healthspan-related gene expression data (describing effects of metabolic intervention associated with improvements in health) onto the aforementioned healthspan pathway maps, we observe the downregulation of Notch signalling in humans and of proliferation/cell-cycle in *C. elegans*. The former reflects the proinflammatory role of the Notch pathway. We identify *transcription*, *proliferation/biosynthesis* and *lipids* as a common theme on the annotation level, and proliferation-related kinases on the gene/protein level. Our literature-based data corpus, including visualization, is available as a reference for future investigations, at http://www.h2020awe.eu/index.php/pathways/.

## Introduction

For a long time, an active, targeted intervention to maintain health into old age was *terra incognita*. It had no priority, and few, if any, reliable data were available to implement it in everyday life. Today, however, systematically established diagnostic hints become available for the individual, based on family history and biomarker data, including genetic variants (polymorphisms). To assess and prevent premature health deterioration successfully, it would therefore be useful (1) to dissect “health” into a set of its most important features, (2) to detail its molecular basis and to map out molecular “healthspan pathways”, and (3) to specify biomarkers and corresponding supportive interventions for the various features of health and for health itself. Arguably, the increase in life expectancy in the last 100 years has not been accompanied by an increase in disease-free life expectancy (Crimmins, 2015), (Robine, Jagger, & Crimmins, 2013). Cardiovascular disease, type-2 diabetes and neurodegenerative disorders are highly prevalent in the elderly, and these disorders frequently coexist in the same aged individual, often with mutual reinforcement (Fillenbaum, Pieper, Cohen, Cornoni-Huntley, & Guralnik, 2000). Extending healthspan may thus enable economic, societal and individual gains on a large scale (Fuellen et al., 2016).

Intervention studies to prolong healthspan based on compound exposure in humans are limited to relatively few compounds. Resveratrol for instance, being one of the best-studied poly-phenols in humans and animals, has been tested in several clinical studies (Berman, Motechin, Wiesenfeld, & Holz, 2017). These include studies focused on biomarkers, like the level of blood glucose (Brasnyó et al., 2011) and cholesterol (S. Chen et al., 2015), or glutathione S-transferase expression (Ghanim et al., 2011). However, data about *long-term* effects on overall health in human are missing in general, and given the average human life expectancy, they are difficult to obtain. Therefore, model organisms are of great relevance to uncover the molecular basis of healthspan and to identify supporting compounds. The nematode *Caenorhabditis elegans (C. elegans)* is a widely used model organism for studying ageing which guided the discovery of fundamental ageing-related findings, e.g., on calorie restriction and Insulin/IGF-1 like signaling (Gruber et al., 2015). Last not least, around 40% of the genes found in *C. elegans* have human homologues (Shaye & Greenwald, 2011) and studies revealing the role of metabolism on health conducted in *C. elegans* have been subsequently strengthened in murine models (Berry & Cirulli, 2013), rendering this nematode a valuable model for human ageing processes. Most recently, *C. elegans* has come to enjoy increasing popularity as a model for health (Sutphin et al., 2017), (Luyten et al., 2016), and an ever increasing number of compounds are tested in *C. elegans* for their anti-ageing and health effects (Chen, Barclay, Burgoyne, & Morgan, 2015), (Luo, Wu, Brown, & Link, 2009), (Collins, Evason, & Kornfeld, 2006), (De Haes et al., 2014).

Here, we assemble and explore “healthspan pathway maps”, that is, annotated sets of interacting genes implicated in health. To create these, we follow a stepwise procedure: first, we dissect health into its various features, with an emphasis on disease and dysfunction. Second, we compile lists of genes associated with health based on the literature, for humans and *C. elegans*. Third, we organize these genes into *maps of healthspan pathways*, based on gene/protein interaction and annotation data. Fourth, we create an overlay of health-related gene expression data onto the resulting healthspan pathway maps, highlighting corroborating knowledge that was not used as input. Finally, we investigate the overlap of the *healthspan pathways* in humans and *C. elegans*.

For humans, we consider that knowledge of the causal basis of health may be best derived from genetic association studies. Based on an extensive review of the literature, we identify a core set of 12 genes that are genetically associated with a lack of frailty (Mekli, Marshall, Nazroo, Vanhoutte, & Pendleton, 2015), (Ho et al., 2011) and the Healthy Aging Index (Sanders et al., 2014), and another set of 40 genes genetically associated with (a lack of) multiple diseases, or with longevity mediated by a lack of disease (see Supplementary Tables 1-3). In contrast to humans, genetic intervention studies on healthspan are available for *C. elegans*, as well as compound intervention data. A lack of dysfunction exemplified by stress resistance, locomotion, pharyngeal pumping and reproduction are taken as the key health features in *C. elegans* (Rollins, Howard, Dobbins, Washburn, & Rogers, 2017). On this basis, a core set of 11 genes is directly implicated in improvements of locomotion by genetics, and another set of 20 genes is indirectly implicated in improvements of the key health features by studies that investigate effects of compounds (see Supplementary Tables 4 and 5).

We then place the genes implicated in health into context by adding gene/protein interaction and gene annotation knowledge. Specifically, we turn the lists of genes into gene/protein interaction networks, to which 20 closely interacting genes are added, employing GeneMania (Zuberi et al., 2013). Gene ontology annotation data are then used to annotate clusters of strongly connected genes within the network, employing AutoAnnotate (Kucera, Isserlin, Arkhangorodsky, & Bader, 2016). Then, we elaborate how the resulting healthspan pathways can be interpreted in plausible ways, specifically in the light of independent health-related gene expression data describing effects of caloric restriction and of rapamycin, and in the light of gene expression data describing aging and disease. We also predict microRNAs that may be potential regulators of healthspan. Finally, we find that if we construct an overlap between the healthspan pathways in *C. elegans* and humans, we find genes involved in transcription, proliferation/biosynthesis and lipids, but this overlap it is not straightforward to interpret in the light of the independent health-related gene expression data that we used to test plausibility of the single-species healthspan pathway maps.

All healthspan pathways discussed in this manuscript, as well as the overlaps we found between species, are available for interactive exploration at http://www.h2020awe.eu/index.php/pathways/.

## Results

### Studies of health

The main sources of knowledge about health, that is, about features, biomarkers and interventions regarding health-related phenotypes, are

a. observational genetic investigations, usually in the form of genome-wide association studies, looking for associations between health and polymorphisms of specific genes,
b. observational studies of non-genetic biomarkers, which are dynamic in time and are usually related to known canonical pathways, and their longitudinal or cross-sectional correlation with health,
c. interventional studies, most often in model organisms, where interventions affecting health may be genetic or based on food or (pharmaceutical) compounds, and the intervention effects are measured on the molecular level, implicating particular genes or pathways.

Like genetic studies, compound intervention studies can, in principle, elucidate the causative basis of health. Studies of type (b) may only be revealing correlative evidence and can sometimes not be linked to particular genes; therefore, we will not consider these further. Candidate biomarkers of health can be many kinds of features with the potential to predict future health better than chronological age (Fuellen et al., 2018); they may be genetic (polymorphisms; such biomarkers are essentially static over lifetime), molecular but not genetic (epigenetic or transcript or protein or metabolic markers, etc.), cellular (blood counts, etc.) or organismic (such as grip strength). Based on studies of types (a) and (c), in this work we will only deal with genes and sets of genes (that is, genes organized into networks or pathways) as candidate biomarkers of health.

### Defining Health

*Health* is a term in biology and medicine that is hard to define. We propose that the best definition of health must be based on an aggregation of the literature, see also (Fuellen et al., 2018), (Luyten et al., 2016). Then, healthspan is simply the time spent in good health. Supplementary Tables 1-3 list features of *human* health as discussed in the literature, referring to lack of dysfunction, lack of *multiple* diseases, and lifespan/longevity mediated by lack of disease. In principle, at least for human, *dysfunction* can be operationalized with the help of a codified classification of function (such as the ICF, the International Classification of Functioning, Disability and Health, www.who.int/classifications/icf/en/). This classification provides criteria to establish that an individual is affected by a dysfunction. As described and discussed in (Fuellen et al., 2018), we can filter the “body function” part of the ICF by looking for follow-up in the literature on health and healthspan. The result is a pragmatic community consensus definition of dysfunction, centering around the lack of physiological, physical, cognitive and reproductive function; a lack of physiological, physical and cognitive functions is often called frailty. To a large degree, this consensus definition can be used for non-human species as well. Further, *disease* can also be operationalized by a codified classification (such as ICD-11, International Statistical Classification of Diseases and Related Health Problems, www.who.int/classifications/icd/en/). Again, the classification provides criteria to establish that an individual is affected by a disease. In this paper, affection by a single disease is not considered, as in old age, single-disease morbidity rarely exists, and in terms of interventions, we are interested in preventing more than one disease. As described and discussed in (Fuellen et al., 2018), not all parts of the ICD feature diseases related to health and healthspan. However, we note that all diseases referred to in Supplementary Tables 1-3 qualify as age-associated diseases.

For *C. elegans*, Supplementary Tables 4-5 list features of health based on the literature, referring to lack of dysfunction in the form of stress resistance (in response to thermal and oxidative stress), (stimulated) locomotion, pharyngeal pumping, and reproduction. These features dominate the literature, and they cover the aspects of physiological function, physical and cognitive function, and reproductive function, as in human (Fuellen et al., 2018). Of note, genetic analyses of health in *C. elegans* have focused up to now mostly on (stimulated) locomotion. Stimulated locomotion integrates some aspects of strength (physical function) and cognition (cognitive function).

### Genes associated with health

Our list of *health genes*, that is, genes with associations to health in humans are based on genetic association, and we can assume some probability of them being causal, leaving aside the intrinsic ambiguities to assign polymorphisms (in the form of SNPs, single-nucleotide polymorphisms) to genes, e.g. in intergenic regions or in intronic regions with overlapping non-coding RNA on the complementary strand (Schwarz et al., 2008). In turn, for *C. elegans*, few studies report effects of genetic interventions on health, though these are increasingly becoming available; (Sutphin et al., 2017) is the only large-scale genetic study that we could identify as of early 2018, even though it is essentially a small-scale study of healthspan based on a large-scale study of lifespan. Many more studies in *C. elegans* refer to canonical aging-related pathways, and in contrast to studies in humans, these studies often directly report the molecular effects of compound intervention. The *C. elegans* genes listed in the supplementary tables are thus based on the effects of genetic intervention and on the effects of compound intervention on the gene level, and we can assume a high probability of causality in both cases.

### Additional genes associated with C. elegans health

For *C. elegans*, we generated an additional list of health-associated genes that cannot be generated for humans (see Experimental Procedures), using *WormBase* to systematically identify health-related compound interventions with associated gene expression data, and compiling the list of genes with strongest differential expression that are well-annotated by Gene Ontology terms.

### From gene lists to maps of healthspan pathways

We used *Cytoscape* and some of its application plugins as the most straightforward tool for obtaining and annotating a connected network of the genes from Supplementary Tables 1-3 and from Supplementary Tables 4 & 5. Specifically, we used *GeneMANIA* to establish a gene/protein interaction network and to add connecting genes, and subsequently we clustered all genes based on their connectivity, and added GO-based annotations, using *AutoAnnotate*. The resulting healthspan pathway maps are presented in the following. Moreover, health-related gene expression data are overlaid onto all healthspan pathway maps and will be discussed as well; these data are describing the effects of caloric restriction (CR) in humans (Mercken et al., 2013) and of rapamycin in *C. elegans* (Calvert et al., 2016), as examples of health-promoting interventions, or they describe the effects of aging and disease in specific tissues.

For humans, the gene list derived from Supplementary Tables 1-3 (see Suppl. Table 6) yielded the network of Figure 1, where the two largest pathways/clusters (15 and 13 genes) are specifically labeled by NOTCH & transcription initiation, and by proliferation, and the smaller pathways/clusters (4, 3, 3 and 3 genes) are labeled by cholesterol & lipid processes, by thymus activation, by myotube (striate muscle) regulation, and by Wnt signalling. In Figure 1 bottom, the list of pathways/clusters is given, and the details of the largest pathway are zoomed in.

**Figure 1:**
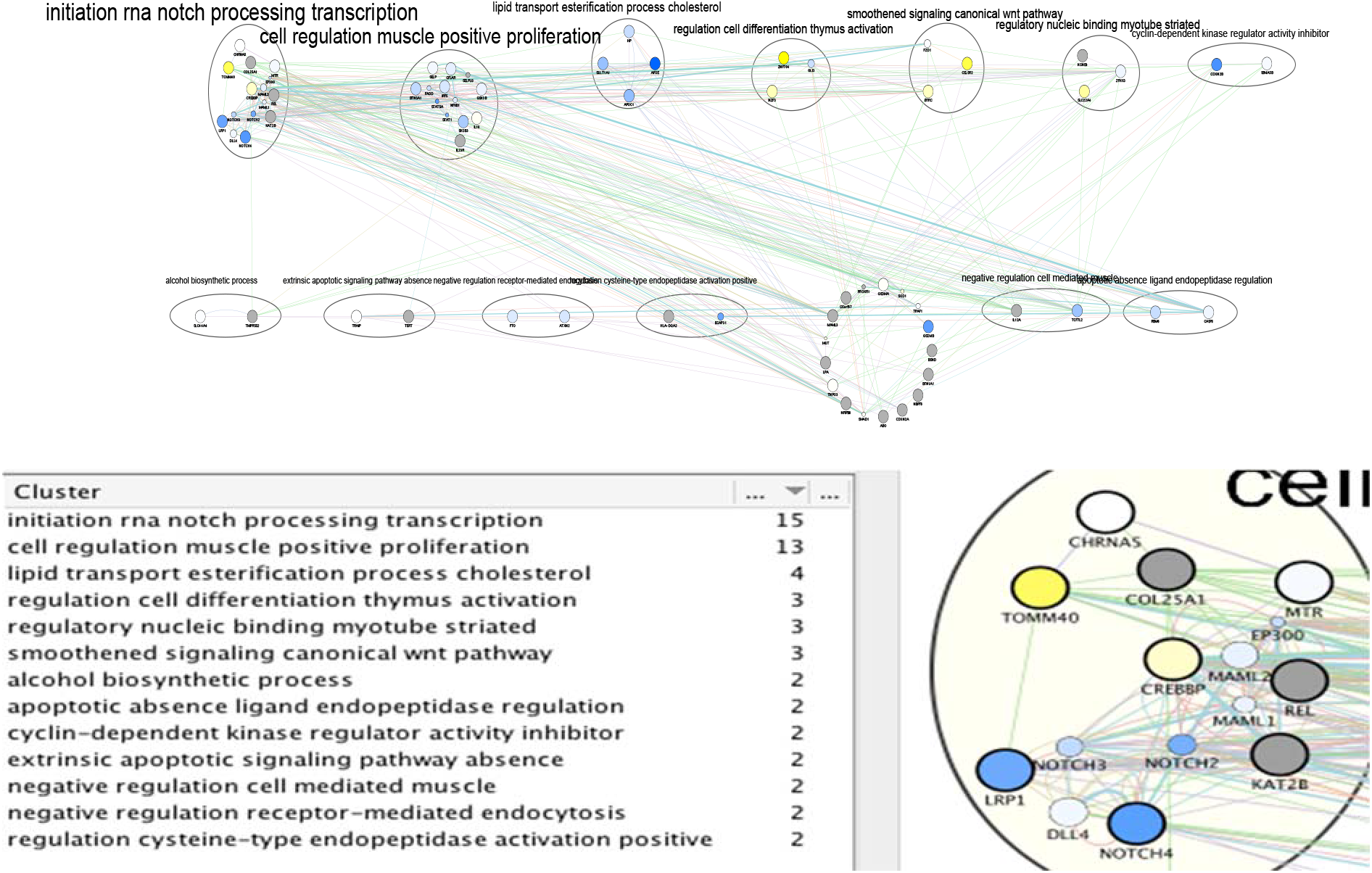
A healthspan pathway map for humans, based on Supplementary Tables 1-3, including the list of pathways/clusters with their labels and size (number of genes), and a zoom into the first pathway. The size of a gene node is proportional to its GeneMANIA score, which indicates the relevance of the gene with respect to the original list of genes to which another 20 genes are added by GeneMANIA, based on the network data. Genes upregulated by CR are shown in yellow, downregulated genes are shown in blue, and grey denotes genes for which no expression values are available in the caloric restriction dataset (Mercken et al. 2013). The color of an edge refers to the source of the edge in the underlying network, that is co-expression (pink), common pathway (green), physical interactions (red), shared protein domains (brown), co-localization (blue), predicted (orange), and genetic interaction (green). The thickness of an edge is proportional to its GeneMANIA “normalized max weight”, based on the network data. Genes from the GeneMANIA input list feature a thick circle, while genes added by GeneMANIA do not.

In the largest pathway/cluster, in light of the CR-triggered gene expression changes, the most prominent findings are an induced downregulation of NOTCH4 (and to a lesser extent of NOTCH 2&3), as well as of LRP1, and an upregulation of TOMM40 and CREBBP (also known as CBP). The family of NOTCH proteins has various functions, including a pro-inflammatory one (Balistreri, Madonna, Melino, & Caruso, 2016), (Zhang, Kuang, Wang, Xu, & Ren, 2017). NOTCH4 is upregulated in kidney failure (Liu et al., 2017), and promotes vascularization/angiogenesis, which includes its upregulation in malignancy (Zhang et al., 2017), (Kofler et al., 2011). A downregulation of NOTCH4 by CR can thus be taken as beneficial effect. This is less obvious for LRP1, the low-density lipoprotein receptor-related protein 1, which is responsible for membrane integrity and membrane cholesterol homeostasis, thus being involved in proper myelination (Lin, Mironova, Shrager, & Giger, 2017) and vascular integrity (Strickland, Au, Cunfer, & Muratoglu, 2014). A downregulation of LRP1 during CR could therefore be seen as deleterious. However, LRP1 expression mainly depends on cholesterol levels (Llorente-Cortes, Otero-Nivas, Sanchez, Rodriguez, & Badimon, 2002) – and these are lower during fasting. Hence, lower LRP1 expression actually reflects a lower LDL level, which per se has been found to be protective, assuming brain levels to follow plasma levels. The upregulations observed for TOMM40 and CREBBP during CR can also be seen as protective. TOMM40 is part of a mitochondrial membrane protein translocase, supporting mitochondrial function (Zeitlow et al., 2017), and low expression and/or particular risk alleles of this protein are associated with Huntington’s and Alzheimer’s Disease (Shirendeb et al., 2011), (Chong et al., 2013). Of note, TOMM40 upregulation during CR goes together with APOE4 downregulation. Although both genes are closely located on chromosome 19, prompting the speculation that this linkage could imply concordant expression changes, this is obviously not the case here. CREBBP is a transcriptional co-activator with histone-acetyltransferase activity (Bedford & Brindle, 2012), acting primarily on histones 3 and 4, and thus it acts in concert with a range of transcription factors. Its downregulation is deleterious, resulting in, e.g., MHCII expression loss on lymphocytes (Hashwah et al., 2017), rendering the lymphocytes dysfunctional for antigen presentation, and in inflammatory signalling (Dixon et al., 2017). An upregulation of CREBBP by CR is thus likely beneficial. We further investigated the miRNAs that are statistically enriched in the largest healthspan pathway using the TFmir webserver (Hamed, Spaniol, Nazarieh, & Helms, 2015), revealing regulation of NOTCH genes implicated in the epithelial-mesenchymal transition, cancer, heart failure and obesity, see Supplementary Results. The genes in the next-largest pathway/clusters, related to cell proliferation and lipids, are also described there in detail, as well as further evidence provided by mapping aging- and disease-related gene expression data onto them, as published or collected by (Aramillo Irizar et al., 2018).

For *C. elegans*, the gene list derived from Supplementary Tables 4-5 (see Suppl. Table 7) yielded the network of Figure 2, where the largest clusters (9 and 6 genes, respectively) are labeled by immune response process and by terms related to the mitochondrion. Three clusters (of 4 genes each) specifically feature dauer/dormancy, regulation and hormone response. In Figure 2 bottom, the list of pathways/clusters, and the details of the largest pathway are zoomed in. Regarding the first pathway, rapamycin induces ets-7 transcription, which was shown to be also necessary for the healthspan-promoting effects of salicylamine (Nguyen et al., 2016). Furthermore, rapamycin upregulates the transcription factor daf-16 (a homolog to Foxo) and downregulates the daf-16 inhibitors akt-1 and akt-2, putatively leading to an improved stress- and immune-response and prolonged lifespan via the Insulin/IGF-1 pathway (Henderson & Johnson, 2001). Along the same lines, the akt-1 and akt-2 activator pdk-1 is also downregulated by rapamycin, further promoting daf-16 activity (Paradis, Ailion, Toker, Thomas, & Ruvkun, 1999). In contrast, the daf-16 inhibitor sgk-1 (a homolog to Nrf) is upregulated; however, its inhibitory role is subject of discussion (Mizunuma, Neumann-Haefelin, Moroz, Li, & Blackwell, 2014). Finally, the transcription factors hsf-1 and skn-1, both important in stress response processes (Morton & Lamitina, 2013); (Wang et al., 2010), are slightly downregulated in rapamycin-treated *C. elegans*. In the Supplementary Results, the next-largest pathway/clusters, related to the mitochondrion, to dauer/dormancy, to regulation, and to hormone response are described in detail.

**Figure 2:**
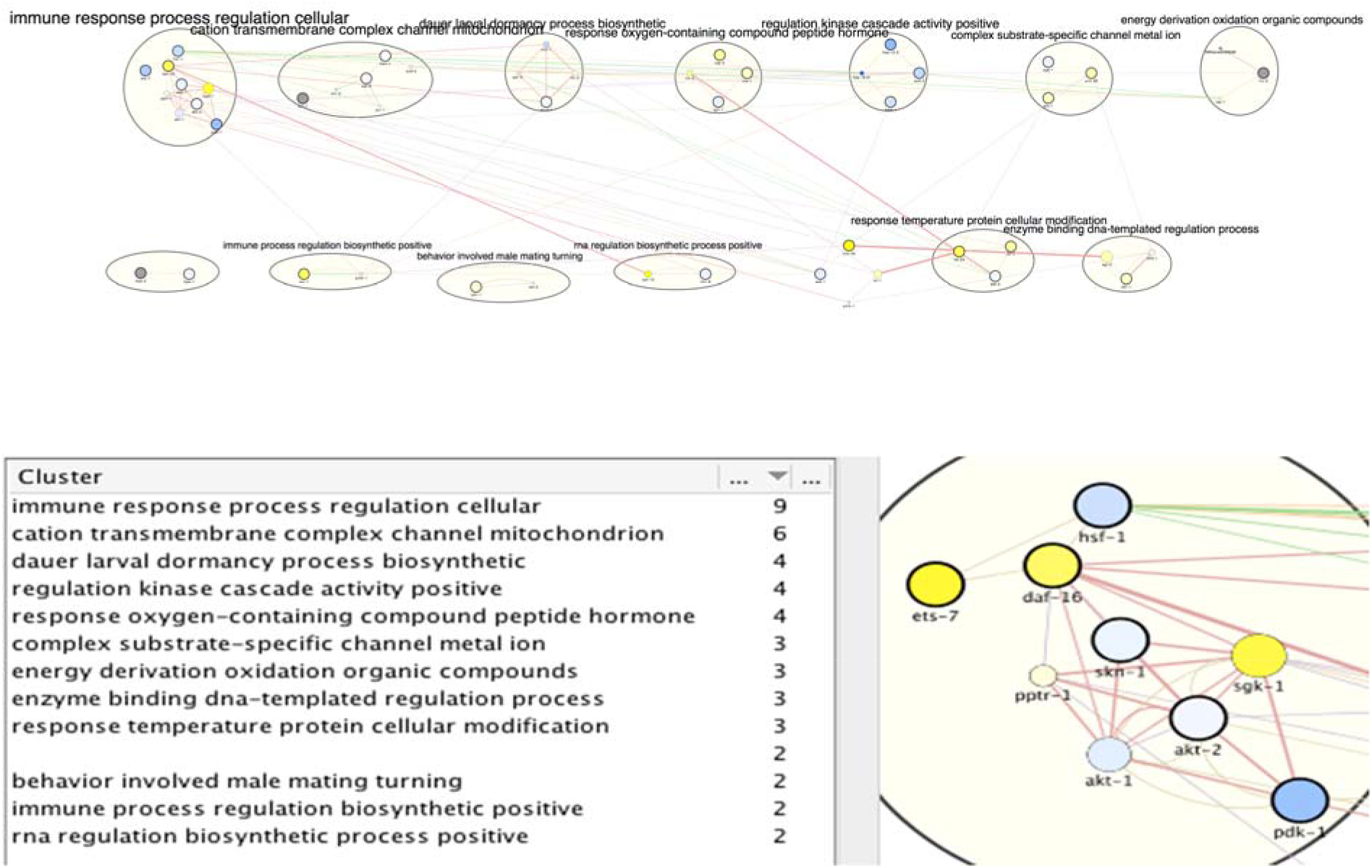
A healthspan pathway map for C. elegans, based on Supplementary Tables 4 and 5. See also Figure 1. Gene expression data reflect the effect of rapamycin (Calvert et al. 2016).

For *C. elegans*, we also derived a gene list from *WormBase*, taking the genes that are most differentially regulated by healthspan-extending interventions and, at the same time, are annotated with a sufficient number of GO terms (see Experimental Procedures; Suppl. Table 8). We obtained the network of Figure 3. Curiously, the top healthspan pathways of 11, 9 and 8 genes are related to the *endoplasmic reticulum (ER*), lipid & membrane, to the *peroxisome*, macrobody & ER, and to the *lysosome*. The endoplasmic reticulum, the peroxisome and the lysosome are part of the endomembrane system, together with the mitochondria, contributing to healthspan and longevity in mammals and beyond (Nisoli & Valerio, 2014). Peroxisomal function connects this pathway to dietary effects on lifespan (Weir et al., 2017), and to liver disease (Cai et al., 2018). The second tier of healthspan pathways (6 or 5 genes) are related to morphogenesis, biosynthesis and transcription.

**Figure 3:**
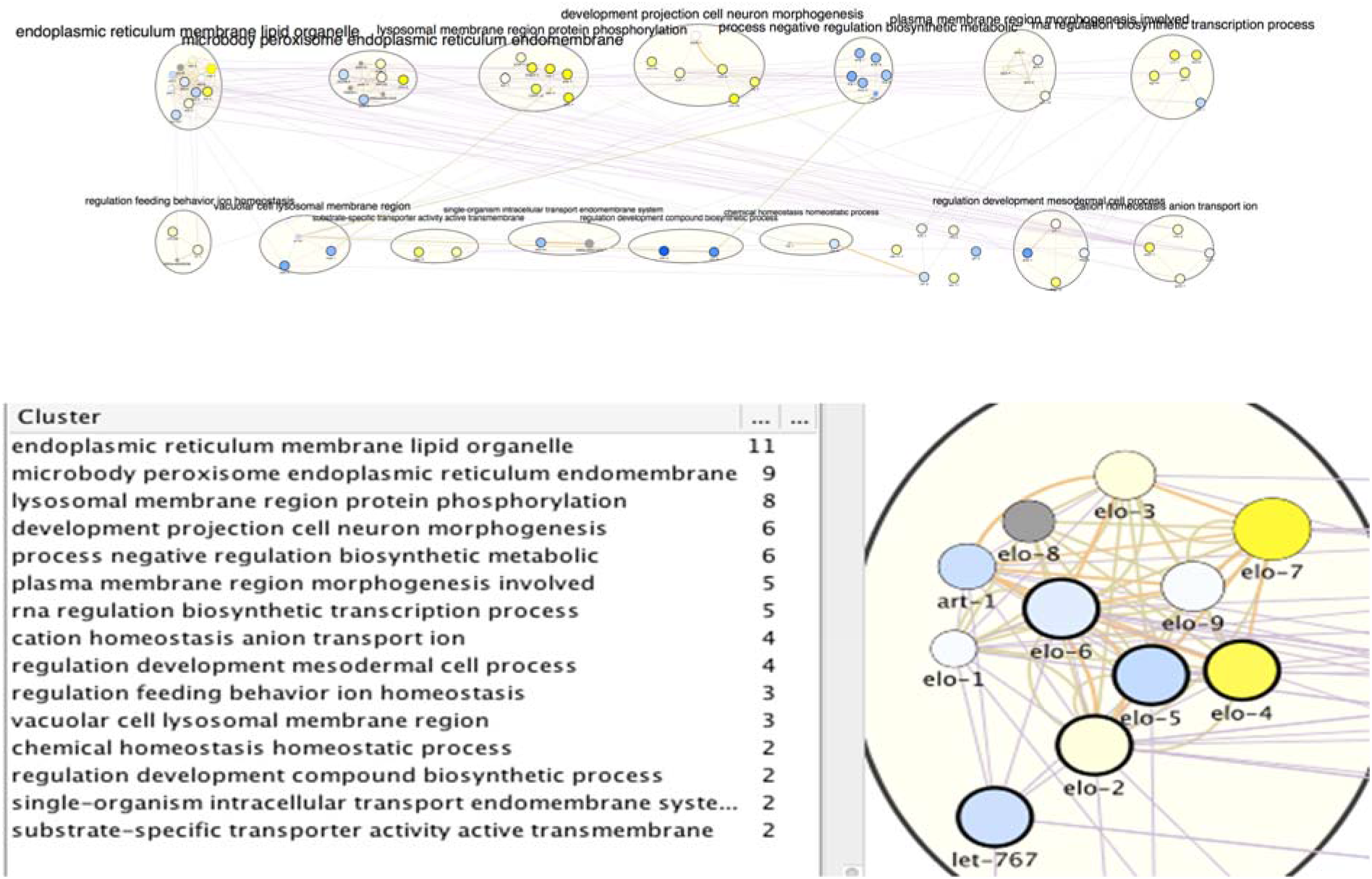
A healthspan pathway map for *C. elegans*, based on genes affected the most by healthspan-extending interventions, using WormBase gene expression data. See also Figures 1 and 2.

For the *WormBase* data, the list of pathways/clusters, and the details of the largest pathway, are given in Figure 3, bottom. The ER/lipid-related pathway includes genes involved in fatty acid elongation/production (elo-1 to elo-9; let-767; art-1). Overlaying the rapamycin gene expression data, the well-characterized elo-1 and let-767 genes show negligible downregulation. However, the importance of elongase genes for health maintenance in general was repeatedly documented. Vásquez and colleagues (Vásquez, Krieg, Lockhead, & Goodman, 2014) demonstrated the impairment of touch response in elo-1 mutants. They argue that elo-1 has a crucial role in the synthesis of C20 polyunsaturated fatty acids which are required for mechanosensation. Moreover, elo-1 mutants showed increased resistance to *Pseudomonas aeruginosa* infections due to the accumulation of gamma-linolenic acid and stearidonic acid (Nandakumar & Tan, 2008) and knockdown of elo-1 or elo-2 extend survival during oxidative stress (Reis et al., 2011). Finally, art-1 is a steroid reductase that is downregulated by rapamycin in our case, but also in long-lived eat-2 mutants (Yuan et al., 2012). In the Supplementary Results, the next-largest pathway/clusters, related to the ER/peroxisome, the lysosome, morphogenesis, biosynthesis and transcription, are described in detail.

### Overlap between human and *C. elegans* health genes and healthspan pathways

Based on reciprocal best orthologs, we found no direct overlap between the human health genes based on genetic associations and the *C. elegans* healthspan genes based in part on genetic interventions, but mostly on expert analysis of intervention effects (Figure 2), or on gene expression changes related to healthspan-extending interventions (Figure 3). We found some hints at an overlap on the level of the healthspan pathway annotations, considering that “proliferation” is listed for human, and “biosynthesis” for *C. elegans*, and “transcription” as well as “lipid” for both.

In search for other modes of overlap, we therefore constructed and compared two interaction networks, based on mapping genes to their respective orthologs in the other species. Each of the two interaction networks is based on the *union set* of the health genes of human (based in turn on genetics, Supplementary Tables 1-3, Figure 1) and of *C. elegans* (based in turn on the gene expression analysis of healthspan-extending interventions using WormBase, Figure 3). Specifically, as outlined in Figure 4, we added the *C. elegans* orthologs of the human health genes to the list of *C. elegans* health genes and *vice versa*, yielding two separate input gene lists for GeneMANIA to enable the construction of the two interaction networks. We used strict ortholog mapping rules (only reciprocal best hits were accepted). By design, the two gene lists feature a high degree of overlap (with differences due to missing orthologs), and their subsequent comparison, consisting of the partial network alignments that are based on ortholog mapping on the one hand and the species-specific network data on the other hand can only reveal *hypotheses for common healthspan pathways*, as long as explicit experimental evidence for a relation to health is only found for one species. Moreover, *interaction points between a healthspan pathway in one species and a healthspan pathway in the other species* may be revealed, if a partial alignment of the interaction networks consists of interacting genes for which the relationship to health was demonstrated only in one species for each pair of orthologs.

**Figure 4:**
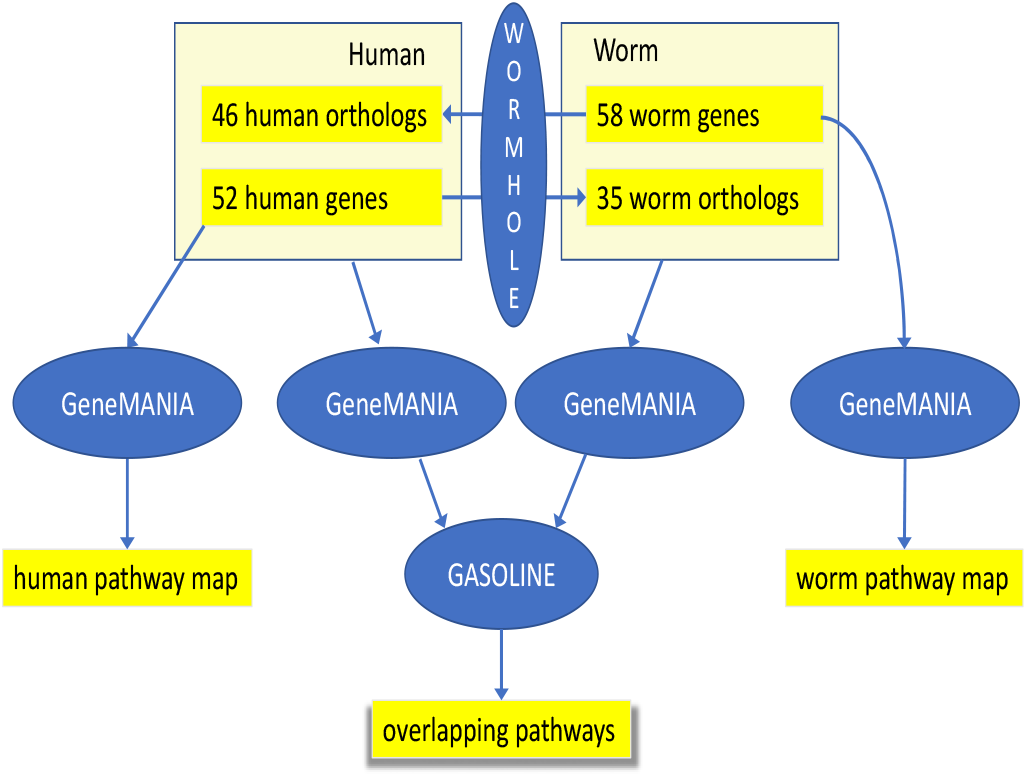
Workflow of the main analysis steps. First, 52 human health genes (Supplementary Tables 1-3) were processed with GeneMANIA and AutoAnnotate to determine the human healthspan pathway map (left, see also Figure 1). Analogously, 58 worm health genes (based on gene expression analysis using Wormbase) were studied, yielding the C. elegans healthspan pathway map (right, see also Figure 3). Then, to determine overlap across species, the gene lists were extended by the orthologs (calculated by WORMHOLE, see Supplemental Experimental Procedures) from the respective other species. We then employed GeneMANIA as before, to generate two interaction networks (one per list). and overlaps between these two networks of health genes were determined by GASOLINE (middle, see also Figure 5).

Of note, of the two interaction networks to be aligned, the first network is based on *C. elegans* health genes, the *C. elegans* orthologs of human health genes, and *C. elegans*gene interaction information provided by GeneMANIA. The second network is based on human health genes, the human orthologs of *C. elegans* health genes, and human gene interaction information provided by GeneMANIA. Despite using similar lists of genes (with differences due to missing orthologs and due to the genes added by GeneMANIA), we can expect that the two GeneMANIA networks are quite different because the interaction data sources employed by GeneMANIA are strongly species-specific. Moreover, we observe that in both cases, the 20 closely interacting genes added by GeneMANIA for one species included no orthologs of the other species. Nevertheless, to identify *joint healthspan pathways* and *interaction points between healthspan pathways*, we used GASOLINE (Micale, Continella, Ferro, Giugno, & Pulvirent, 2014) to align the two networks wherever feasible, obtaining two partial (subnetwork) alignments as output, as shown in Figure 5.

**Figure 5:**
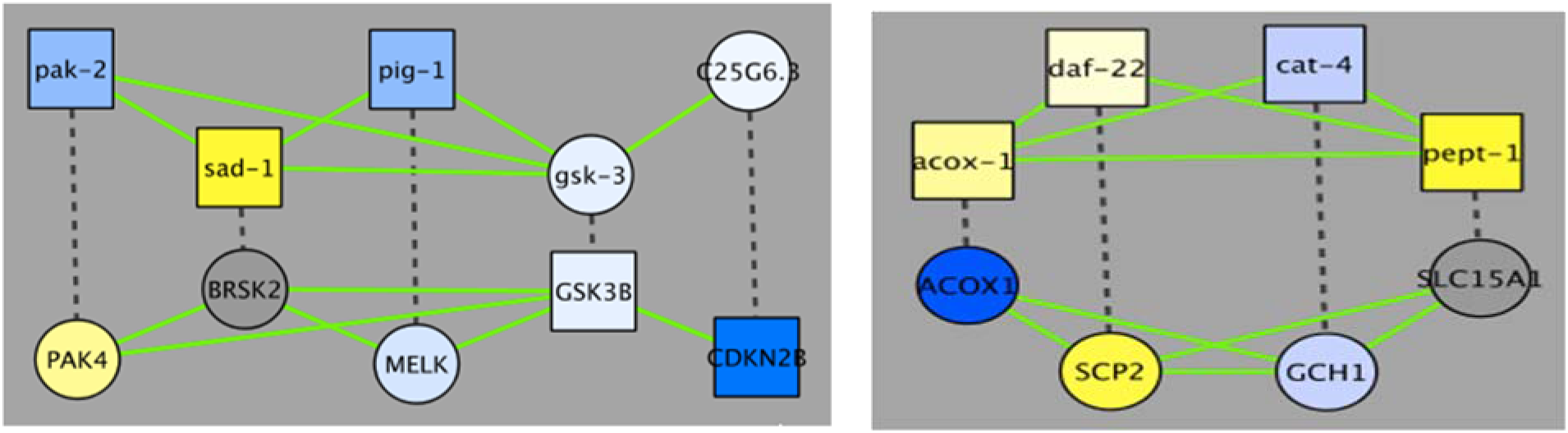
The two alignments demonstrating overlap of (putative) healthspan pathways in human and *C. elegans*, based on a GASOLINE alignment of the network of genes implicated in health-related gene expression changes in WormBase (top), and in human health based on genetic studies (bottom), and of corresponding orthologs. Dashed edges indicate orthologs, green edges indicate interactions based on GeneMANIA known for the respective species; the node shape is square if the gene originates from the original lists of health genes and it is circular if the gene is an ortholog, and node colors are based on gene expression changes triggered by rapamycin (in case of *C. elegans*) or by caloric restriction (in case of human), as in Figures 1–3.

In the first alignment (Figure 5, left), we see an alternating pattern of demonstrated health-relatedness, since pak-2, sad-1 and pig-1 are considered health-related by gene expression analysis using WormBase, while CDKN2B and GSK3B are known to be human health genes (Supplementary Tables 1-3; GSK3B was implicated by a GWAS of the Healthy Aging Index, while CDKN2B was in fact one of the few genes implicated by two independent health studies). The *C. elegans* genes belong to three small clusters in the healthspan pathway map of Figure 3 (pak-2: lysosomal, sad-1: neural, pig-1: biosynthesis), while the human genes belong to one large (GSK3B: proliferation) and one small (CDKN3B: cyclin-dependent kinase) cluster in the human healthspan pathway map of Figure 1. Interactions in *C. elegans* are all based on shared domains (kinase signaling, except for the predicted interaction of gsk-3 & C25G6.3, which is based on the Interologous Interaction Database), while interactions in human are based on shared domains, genetic interaction (i.e., large-scale radiation hybrid) and pathway data. Essentially, the healthspan pathway overlap suggested by our analysis involves serine/tyrosine kinase signalling (pak-2/sad-1/pig-1 & PAK4/BRSK2/MELK), Wnt signalling (GSK3) and cyclin-dependent kinase signalling (CDKN2B). Both alignments are described further in detail in the Supplementary Results.

## Discussion

### Healthspan pathway mapping, current possibilities and future opportunities

We took a pragmatic approach to investigate the molecular basis of health. For humans, there are only few alternatives to genetic association studies, to determine the genes responsible for the phenotype of lacking disease and dysfunction. Post-mortem analyses aside, studies in human are bound to the insights gathered from the etiology and treatment of diseases in patients. Genetic manipulation is possible in human cell lines, but this requires to break down the disease or dysfunction to a cellular phenotype. On the other hand, molecular insights can be gained with few ethical constraints in non-vertebrates such as *C. elegans*. A range of age-associated phenotypes can be observed in *C. elegans* that are similar to human, yet the nematode is still a much simpler organism with far fewer cells and a smaller genome. As noted, however, sequence similarity supports the mapping of ortholog genes across species, suggesting functional similarity or equivalence, and many core pathways are preserved. We discuss the biological interpretation of the lack of evolutionary conservation in the Supplementary Discussion.

Looking for models of human health, there is a choice of cell lines, organoids (“organ-in-a-dish”), *in silico* simulations and model organisms. A major disadvantage of cell lines is their artifactually isolated cultivation. This disadvantage is addressed by organoid-based approaches, which, however, are still far from the human situation they are supposed to model, in terms of size, complexity, and the duration of the investigation. *In silico* simulations are limited in complexity, and their faithfulness may be questioned on many accounts. This faithfulness is also a big issue with model organisms, where ease of handling and a reasonable duration of the investigation usually result in difficult tradeoffs with faithfulness. A discussion of the various approaches just mentioned, and of the various choices of model organisms is beyond the scope of this paper. But it should be mentioned that while health can be defined and investigated in *C. elegans*, this model is also fraught with an array of issues, some shared with organoid-based approaches, but also including some particularities of the species (post-mitotic tissues, artificial cultivation, hermaphrodite, genetic uniformity of strains, lack of organs present in humans such as a heart) and of our modes of investigation (limitations of the data currently available, limitations of healthspan assays in terms of their design and the controlled environment in which these are conducted). Nevertheless, for studies in a living system to be completed in a short time-frame, *C. elegans* is arguably the best choice.

Given lists of genes, there is a plethora of possibilities to organize the genes into groups of related ones. Motivated by the idea of a “healthspan pathway”, we hypothesized that the genes should be known to interact based on functional gene/protein interaction data (provided by GeneMANIA). Here, as in most other studies, pathways are not assumed to be linear. The (higher-level) interaction among the clusters/healthspan pathways (i.e., the pathway map) is given by the individual gene/protein interactions that are shown *between* the clusters in Figures 1-3. However, we did not investigate these further.

The small amount of healthspan gene/pathway overlap that we found may be seen from a pessimistic or an optimistic perspective, depending in part on expectations. From the pessimistic perspective, the molecular processes may be completely different, and the*C. elegans* orthologs of the human health genes are involved in different processes as compared to the human health genes, and *vice versa*. From the optimistic perspective, it may just be that the number and scope of the investigations that yielded the health genes we studied is still insufficient, annotations are still incomplete, and considering only reciprocal best orthologs may be too restrictive. (We tried a less restrictive mapping of orthologs by relaxing the condition that orthologs must be reciprocal, but the overlap was still negligible; results not shown). Nevertheless, future genetic studies are expected to yield more health genes in both species, and their characterizations are expected to improve. Moreover, when we analyze in detail the effects of intervention studies in *C. elegans*, we do find clear hints to some mechanisms that underlie healthspan also in human. For example, changes in the Ins/IGF-1 pathway genes daf-2 and daf-16 are found to be associated with many of the features described in Supplementary Table 5, suggesting a fundamental role for immune defense mechanisms (and proliferation) in health maintenance, as described by (Ermolaeva & Schumacher, 2014).

Of course, the precise definition of phenotype is crucial. If the samples are not really about (lack of) health, in human or in *C. elegans*, then any subsequent molecular or bioinformatics analyses will compare apples and oranges and may thus fail. Therefore, it is important to use a good phenotyping of health in human as well as in *C. elegans*,and on this basis, to collect data as genome-wide as possible. It is evident that there is no *C. elegans* counterpart to most of the (age-related) diseases that we use to define health in humans, and that the aging process that may underlie most of these age-related diseases is poorly characterized and hard to quantify in humans. Nonetheless, locomotion degrades with age in both species, due to changes at the muscle as well as neural level. Two related features of physical function, that is, grip strength (Leong et al., 2015) and the ability to sit and rise from the floor (Brito et al., 2014) are good predictors of all-cause mortality in humans. Likewise, both in humans and in *C. elegans*, the ability to withstand various forms of stress decreases with age (Dues et al., 2016), (Bajorat et al., 2014). Thus, at the level of organs or functional systems, both *C. elegans* and humans show age-related declines in performance, that may well be due to underlying processes that are similar at the cellular and molecular level. We claim that within the limitations of currently available data, the health genes we assembled, the healthspan pathways we constructed based on these, and the overlap we then found between species, are a first glimpse of the species-specific and crossspecies molecular basis of health.

### Experimental Procedures

#### Gene sets associated with health, literature-based

In this work, we conducted a semi-systematic review, using *health*, *healthspan* and *healthy aging*, for human and *C. elegans*, as search terms in *Google Scholar*, initially filtering for recent reviews and considering only the top hits. For humans, genetic studies of Supplementary Tables 1-3 are often not found using health-related keywords, so we included terms related to dysfunction (such as *frailty*) and disease (such as *multi-morbidity*) as well. For the genetics of human frailty, we identified two publications (Mekli et al., 2015), (Ho et al., 2011). Overall, a list of 52 genes (Supplementary Table 1, 12 genes; Supplementary Tables 2 and 3, 40 genes) was taken as the starting point in humans. For the genetics of *C. elegans* health, we followed a similar approach (Supplementary Table 4). For compound interventions in *C. elegans*, we identified a specific set of recent reviews (see Supplementary Table 5). Overall, a list of 31 genes (Supplementary Table 4, 11 genes; Supplementary Table 5, 20 genes) was taken as the starting point in *C. elegans*. From the original publications and reviews, we extracted the gene names, using *ihop* (Hoffmann & Valencia, 2004) to assign HUGO nomenclature names if necessary. In the Supplement, we describe in detail how a second set of health-associated genes in *C. elegans* was identified using WormBase.

#### Construction of maps of clusters/pathways

For all gene sets analyzed, we used the Cytoscape 3.5.1 application *GeneMANIA* (Zuberi et al., 2013), version 3.4.1, downloaded October 2017, with default settings, to create a functional interaction network that is complemented with the GeneMANIA default of 20 connecting genes. For clustering, and for annotating the clusters based on the “annotation name” column of GO annotations collected by GeneMANIA, we used *AutoAnnotate* (Kucera et al., 2016) v1.2, downloaded October 2017, in Quick start modus to enable to “layout network to prevent cluster overlap”, so that a map of disjoint clusters (healthspan pathways) was generated, supplemented by a second advanced annotation step to increase the “max. number of words per cluster label” to the largest possible value of 5. Cluster annotations were generated using WordCloud (Oesper, Merico, Isserlin, & Bader, 2011) v3.1.1, downloaded January 2018.

In the Supplement, we further describe in detail how we overlaid expression data onto the pathway maps, constructed the overlap of healthspan pathways in *C.elegans* and humans, and programmed the web presentation.

## Acknowledgements

We thank Yasmeen Quawasmeh for technical assistance. We acknowledge assistance by Giovanni Micale in using GASOLINE. This project has received funding from the European Union’s Horizon 2020 research and innovation programme under Grant agreement No 633589 (Aging with Elegans). This publication reflects only the authors’ views and the Commission is not responsible for any use that may be made of the information it contains. AAC is supported by a CIHR New Investigator Salary Award and is a member of the Fonds de recherche du Québec – Santé funded Centre de recherche du CHUS and Centre de re-cherche sur le vieillissement.

## Authors’ contributions

Study design: GF, WL, SM, NS. Collection of data: GF, NS, SM. Analysis of data: GF, NS, RK, SSe, HME, CJ, SS, IB, MH. Website: SM, PAd, JV. Manuscript writing: GF, NS, SM, AAC, RK, SSe, HME, LS, BW, FC, AB, PAn, HJG, DR, MB, LJ. All authors reviewed and approved the final manuscript.

## Supporting Information listing

Supplementary Table 1. Features of human health, lack of dysfunction(s), and associated genes.

Supplementary Table 2. Features of human health, (lack of) multiple diseases, and associated genes.

Supplementary Table 3. Features of human health, lifespan/longevity mediated by lack of disease, and associated genes.

Supplementary Table 4. Features of *C. elegans* health, based on genetic studies of health, and associated genes.

Supplementary Table 5. Features of *C. elegans* health, based on compound intervention studies affecting health, and associated genes.

Supplementary Table 6. Human genes associated with health, listing all genes from Tables 1+3.

Supplementary Table 7. *C. elegans* genes associated with health, listing all genes from Tables 4+5.

Supplementary Table 8. *C. elegans* genes associated with health, listing genes based on WormBase gene expression data.

Supplementary Table 9: List of diseases and functional terms associated with the miRNAs enriched as regulators of the largest human healthspan pathway.

Supplementary Table 10: List of diseases associated with the miRNAs enriched as regulators of the second-largest human healthspan pathway.

Supplementary Figure 1: The regulatory interactions between the largest human healthspan pathway and the corresponding enriched miRNAs.

Supplementary Figure 2: The regulatory interactions between the second-largest human healthspan pathway and the corresponding enriched miRNAs.

Supplementary Figure 3: Comparison of expression patterns in two aging tissues, largest human healthspan pathway.

Supplementary Figure 4: Comparison of expression patterns in two aging tissues, second-largest human healthspan pathway.

Supplementary Figure 5: Comparison of expression patterns in two aging tissues, third healthspan pathway.

Supplementary Figure 6: Comparison of expression patterns in three disease-affected tissues, top 3 healthspan pathways.

Supplementary References

